# Sampling strategies to assess microbial diversity of Antarctic cryptoendolithic communities

**DOI:** 10.1101/676775

**Authors:** Claudia Coleine, Jason E. Stajich, Nuttapon Pombubpa, Laura Zucconi, Silvano Onofri, Laura Selbmann

## Abstract

Describing the total biodiversity of an environmental metacommunity is challenging due to the presence of cryptic and rare species and incompletely described taxonomy. How many samples to collect is a common issue faces ecologists when designing fieldwork sampling: collecting many samples may indeed capture the whole metacommunity structure, but can be prohibitively costly and lead to an enormous amount of data to analyse. Conversely, too few samples may yield inadequate and incomplete data which can prohibit complete assessment of community diversity. High-throughput sequencing allows examination of large numbers of samples enabling comprehensive biodiversity assessments. In this study, we sought to estimate how the scale of sampling affects accuracy of community diversity description in order to develop strategies to exhaustively describe the microbial diversity of cryptoendolithic communities in the McMurdo Dry Valleys in Antarctica accounted as the closest Martian analogue on Earth, exhibiting extreme conditions such as low temperatures, wide thermal fluctuations, low nutrient availability and high UV radiation. We found that sampling effort, based on accumulation curves analysis, had a considerable impact on assessing species richness and composition in these ecosystems, confirming that a sampling as large as nine rock specimens was necessary to detect almost all fungal species present, but was not sufficient to capture whole bacterial assemblage.

## Introduction

How much sampling is appropriate to assess the microbial diversity? The number of samples to collect is an important consideration for experimental design of microbial community studies. Large numbers of samples may be required to test ecological hypotheses that depend on accurately capturing the entire metacommunity (Bonar *et al*., 2011), but these requirements may be prohibitively expensive, but too little sampling may yield inadequate data. Therefore, a cost-benefit analysis to maximize data quality and completeness (Gotelli and Colwell, 2001) must be incorporated when designing a field campaign while considering minimal required sampling efforts to achieve the goals (Kasprzak, 1993).

Environmental samples typically contain many rare (low-abundant) taxa or species potentially difficult to identify (Grey et al., 2018). High-throughput sequencing (HTS), particularly environmental metabarcoding (eDNA), have provided new opportunities to analyse large numbers of sample for low costs (Caporaso et al., 2012), leading to a more accurate exploration of biodiversity and community’s composition (see Buée et al., 2009; Jumpponen and Jones, 2009; Blaalid et al., 2012).

Accumulation curves are used to assess if representative microbial diversity is fully sampled with an HTS approach. These assessments may be used to test if sufficient sampling efforts have been performed to accurately estimate the total species observed (Gotelli and Colwell, 2001). Theoretically, finding out how many species characterize a microbial assemblage means sampling more and more individuals until no additional species are found, but often the number of individuals that must be sampled to reach this result may often be prohibitively large (Chao et al., 2009). Exhaustive description of biodiversity meaning detection of all organisms in a community is challenging (Ariño et al., 2008): a sampling plan must be feasible and cost effective, answering a series of biological questions and leading to the best results and proper interpretation of the analysis. Development of a clear community research questions is the key to designing a good sampling plan and more necessary in the extremes and in difficult to access ecosystems such as Polar regions.

The McMurdo Dry Valleys in Southern Victoria Land (Continental Antarctica) are considered one of the most extreme environment on Earth; for the different stressors characterizing this area, such as extreme temperature fluctuations (Doran et al., 2002), dryness, oligotrophy and high levels of solar and UV radiation (Wynn-Williams, 1990; Smith et al., 1992), it is accounted as the closest Martian analogue on Earth (Horowitz et al., 1972; McKay, 1993; Wynn-Williams and Edwards, 2000; Onofri et al., 2004; Baqué et al., 2013). In these ice-free areas, the prohibitive factors limit most life forms and microorganisms can only find refuge in cryptic niches, developing within airspaces of porous rocks and forming cryptoendolithic communities (Friedmann and Ocampo, 1976; Friedmann, 1982; 1988); the endolithic lifestyle is considered a stress avoidance strategy (Pointing and Belnap, 2012; Pointing et al., 2014), providing microorganisms a means to survive in extreme aridity.

Despite recent molecular studies have contributed provided new insight into the biodiversity and distribution of Antarctic cryptoendolithic communities and their response to environmental pressure (de La Torre et al., 2003; Wei et al., 2016; Archer et al., 2017; Selbmann et al., 2017; Coleine et al., 2018a, b), the sampling effort necessary for a proper whole metacommunity description remain still unexplored. In this survey, we aimed to determine the minimum sampling effort required to sufficiently estimate the scale of microbial biodiversity and composition of the Antarctic cryptoendolithic communities in McMurdo Dry Valleys, based on fungal and bacterial metabarcoding profiles. Our findings provide recommendations for optimizing future experimental sampling design for surveys of these communities. These sampling suggestions are critical to reduce possible impact in these strictly protected and sensitive environmental areas (ASMA, Antarctic Specially Managed Areas, and ASPA, Antarctic Specially Protected Areas), with very limited access.

The results include an assessment of the variability of community structure in a sampling of as many as nine rock samples collected from a single locality of the McMurdo Dry Valleys and an estimate of minimum sampling efforts required to adequately characterize fungal and bacterial diversity.

## Materials and Methods

### Sampling area

Nine homogenous colonized by lichen-dominated cryptoendolithic communities sandstones were collected by L. Selbmann and L. Zucconi during the XXVI Italian Antarctic Expedition (December 2010-January 2011) at Battleship Promontory 76°54’36’’S 160°56’05’’E (McMurdo Dry Valleys, Southern Victoria Land, Antarctica) (Fig.1) at 1000 m elevation and 33.5 km distance from sea within an area of about 100 m^2^. The presence of endolithic colonization was assessed by direct observation *in situ*. Rocks were excised using a geological hammer placed in sterile bags and stored at −20°C until processing to extract DNA and perform molecular analysis at University of Tuscia, Italy.

### DNA extraction and metabarcoding sequencing

Sandstones were crushed under sterile conditions to collect 1 g of powdered rock used for DNA extraction with MOBIO Power Soil DNA Extraction kit (MOBIO Laboratories, Carlsbad, CA, USA) following the manufacturer’s instructions. The Ribosomal Internal Transcribed Sequence 1 region (ITS1) and V4 region of the 16S rRNA gene were targeted to assess the fungal and bacterial community membership, respectively. We amplified the ITS1 region using barcoded primers ITS1F/ITS2, developed for shorter read length (Smith and Peay, 2014) and V4 region using the new developed barcoded F515/R806 primer set as described by Caporaso and colleagues (2012) (Table 1). PCR was carried out with a total volume of 25 μl, containing 1 μl of each primer, 12.5 μl of Taq DNA Polymerase (Thermo Fischer Scientific Inc., Waltham, MA, USA), 9.5 μl of nuclease-free water (Sigma–Aldrich, UK) and 5 ng of DNA template using an automated thermal cycler (BioRad, Hercules, CA). The ITS1 locus was amplified following initial denaturation at 94°C for 1 min, 35 cycles of denaturation at 94°C for 30 s, annealing at 52°C for 30 s, extension at 68°C for 90 s, followed by a final extension at 68°C for 7 min. The PCR for the V4 region followed a protocol of an initial denaturation at 94°C for 3 min, 35 cycles of denaturation at 94°C for 45 s, annealing at 50°C for 1 min, extension at 72°C for 90 s, followed by a final extension at 72°C for 10 min. Amplicons, quantified by Qubit dsDNA HS Assay Kit (Life Technologies, USA), were pooled and purified with Qiagen PCR CleanUp kit (Macherey-Nagel, Hoerdt, France). Paired-end sequencing (2×300 bp) was carried out on Illumina MiSeq platform at the Institute for Integrative Genome Biology, University of California, Riverside, Riverside, California.

DNA extraction was performed in duplicate and replicates were sequenced separately.

**Table 1.**
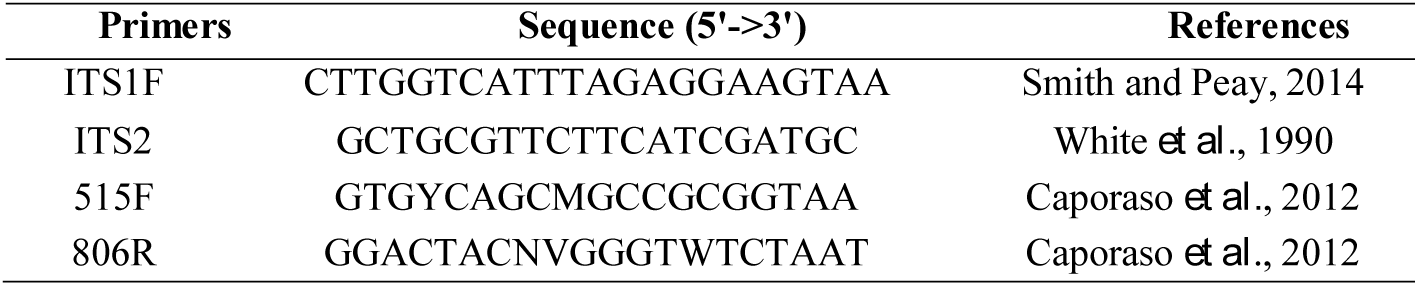
ITS and 16S primer sequences for metabarcoding sequences.

### Processing of metabarcoding data

The ITS1 and 16S datasets were processed separately with the AMPtk (Palmer et al., 2018; https://github.com/nextgenusfs/amptk) tool. Briefly, overlapping 2×300 base pair MiSeq reads were merged using USEARCH v8 (Edgar and Flyvbjerg, 2015). Barcodes and primers were removed from the amplicons sequencing data and reads were demultiplexed using the script split_libraries.py in QIIME v1.9.1 (Caporaso et al., 2010). Reads were then subjected to quality trimming based on accumulation of expected errors less than 1.0 (Edgar and Flyvbjerg, 2015) and chimeric reads were removed in AMPtk as implemented by USEARCH (Edgar, 2010). A 97% sequence similarity clustering threshold was utilized to create Operational Taxonomic Units (OTUs) using VSEARCH (v 2.3.2) (Rognes et al., 2016) tool. Taxonomy assignment was performed with SINTAX/UTAX (Edgar, 2010) (Supplementary Materials Tables S2, S3)

Singletons (OTUs represented by 1 reads in whole dataset) and rare taxa (<5 reads) were discarded as likely false positives due to sequencing errors, as recommended by Lindahl et al. (2013). Reads obtained from duplicates were merged. Fungal and bacterial rarefaction curves were calculated using the ‘rarecurve’ function in the package ‘vegan’ (v. 2.3-4; Oksanen et al., 2013) in R 3.2.0 (R-Development-Core-Team 2015).

Raw sequence data were submitted to the GenBank database under BioProject accession number PRJNA379160.

### Biodiversity and NMDS ordination plots

Alpha diversity from the OTU tables was estimated using Primer-E v7 (Auckland, New Zealand) software, calculating biodiversity indices as follows: (i) the species richness (S) as a count of the total number of OTUs found in each sample; (ii) the Shannon index (H’), a phylotype-based approach constructed using OTU groupings (Shannon and Weaver, 1949; Ludwig, 1988); (iii) the Simpson’s Index of Dominance (1-D), measuring the probability that two individuals randomly selected from a sample will belong to the same species (Simpson, 1949). Statistical analysis was performed by one-way analysis of variance (one-way ANOVA) and pairwise multiple comparison procedure (Tukey test) carried out using the statistical software SigmaStat 2.0 (Jandel, USA).

Changes in fungal and bacterial assemblages’ composition were evaluated with non-metric multidimensional scaling (NMDS) analysis, performed with PAST (v.2.17) (PAleontological Statistics) (Hammer et al., 2001) and their confidence intervals were investigated through permutational PERMANOVA (Anderson et al., 2013). We used Jaccard (presence–absence) and Bray–Curtis (relative abundance) metrics to calculate pairwise community distance matrices and examine differences in beta diversity. Abundance data were square-root transformed to limit the influence of OTUs with high sequence counts and analyses were carried out with 999 permutations. A small probability p-value (<0.05) indicated a significant difference of community composition among all samples.

### Accumulation curves and extrapolation analysis

Species accumulation curves analysis was implemented to compare microbial assemblages and to estimate the expected number of new species given additional sampling e□ort, which can give insights to proper sampling protocols (Colwell and Coddington, 1994; Moreno and Hal□ter, 2000).

Curves were calculated using PRIMER-E v7 implementing Plots>Species Accum Plot function. The increasing total number of different species observed (S) was plotted against samples successively pooled (referred to as the Sobs curve; Sobs: total number of species observed in all samples pooled). We used the ‘permute’ option to enter samples utilizing randomization procedures (Colwell and Coddington, 1994; Gotelli and Entsminger, 2001). When the curve reaches an asymptote, and the number of OTUs does not increase by adding further samples, it indicates sufficient samples have been collected to accurately characterize the community (Pielou, 1977).

We also computed an extrapolation model (sample-based extrapolation curve) with 95% unconditional confidence intervals, using the analytical formulas of Colwell et al. (2012). The model attempts to predict the asymptotic number of species that would be found for an increasing number of samples, estimating the number of samples needed to reach a plateau, by using EstimateS v9 (Statistical Estimation of Species Richness and Shared Species from Samples) (Colwell et al., 2012).

For all tests, sample-based incidence data were used and analysis carried out with 999 permutations.

### Quantitative PCR

Quantification of fungal and bacterial rRNA gene copies was estimated by using quantitative PCR assay. To estimate fungal and bacterial gene abundances, standard curves were generated using a 10-fold serial dilution of a plasmid containing a copy of *Cryomyces antarcticus* ITS rRNA gene or *Escherichia coli* 16S rRNA gene. Amplicons were generated in 100 μl reactions, containing 50 μl of 2x PCR BioMix™ (Bioline, London, UK), 1 ul (5 pmol) of both forward and reverse primers, 43 μl of DEPC water and 5 μl template of genomic DNA. For fungi, the primers used were NS91F (5’-GTCCCTGCCCTTTGTACACAC-3’) and ITS51R(5’-ACCTTGTTACGACTTTTACTTCCTC-3’), while Eub338 (5’-ACTCCTACGGGAGGCAGCAG-3’) Eub518 (5’-ATTACCGCGGCTGCTGG-3’) primer set was utilized for bacterial component (Fierer et al., 2005). Amplification of ITS region was carried out as follows: 95 °C for 3 min and the 45 cycles of 95 °C 40 s, 55 °C 30 s and final extension at 72 °C 40 s; while 16S region was amplified as follows: 95 °C for 3 min and the 45 cycles of 95 °C 40 s, 53 °C 30 s and final extension at 72 °C 40 s. PCR products were purified using the NucleoSpin® PCR Clean-up kit (MACHEREY-NAGEL, GmbH & Co. KG), quantified with Qubit dsDNA HS Assay kit and then cloned using the pGEM®-T Easy Vector Systems (Promega, Madison, Wisconsin, US). Plasmids were isolated using the NucleoSpin Plasmid kit (Macherey-Nagel, GmbH & Co. KG). Five standards were utilized for qPCR in series from 10^3^ to 10^7^ copies.

The 25□μl qPCR reactions contained 12.5□μl iQ™ SYBR® Green Supermix (Bio-Rad, Hercules, California, US), 1□μl of each pM forward and reverse primers, 0.3 ng of environmental DNA or standard and 9.5□μl nuclease-free water. The reactions were carried out on a Quantitative real-time BioRad CFX96™ PCR detection system (Bio-Rad, Hercules, California, US). Primers were the same used to generate standard curves and the PCR protocols were those mentioned above.

Melting curve and gel electrophoresis analyses confirmed that the amplified products were of the appropriate size. Fungal and bacterial gene copy numbers were generated using a regression equation for each assay relating the cycle threshold (*C*_t_) value to the known number of copies in the standards.

All qPCR reactions were run in triplicate.

Fungal-to bacterial (F/B) ratio was also calculated based on log-transformed abundance values. Means and standard deviations were calculated and statistical analysis were performed using one-way analysis of variance (ANOVA) and pairwise multiple comparison procedures, carried out using the statistical software SigmaStat v2.0 (Jandel, USA). Significant differences were calculated by Tukey test with p value<0.05.

## Results

### Metabarcoding analysis

Fungal and bacterial reads were retrieved from all samples. For ITS 558,579 reads passed quality filtering and the per-sample number of rRNA gene reads averaged 62,064 with a minimum of 36,246 to a maximum of 86,668 reads. A total of 624,195 16S sequences reads were generated across the samples with an average of 69,355 reads, minimum of 49,597 and maximum 111,307 reads per sample. After removal of global singletons and rare taxa (<5 reads) (56 out of 245 fungal OTUs and 198 out of 715 bacterial OTUs).

Sequence reads were grouped, using a 97% identity threshold, into 189 fungal and 517 bacterial OTUs, respectively; the number of reads for each sample is reported in Table 2. Rarefaction curves were constructed and results are reported in Figs. 1S, 2S.

**Table 2.**
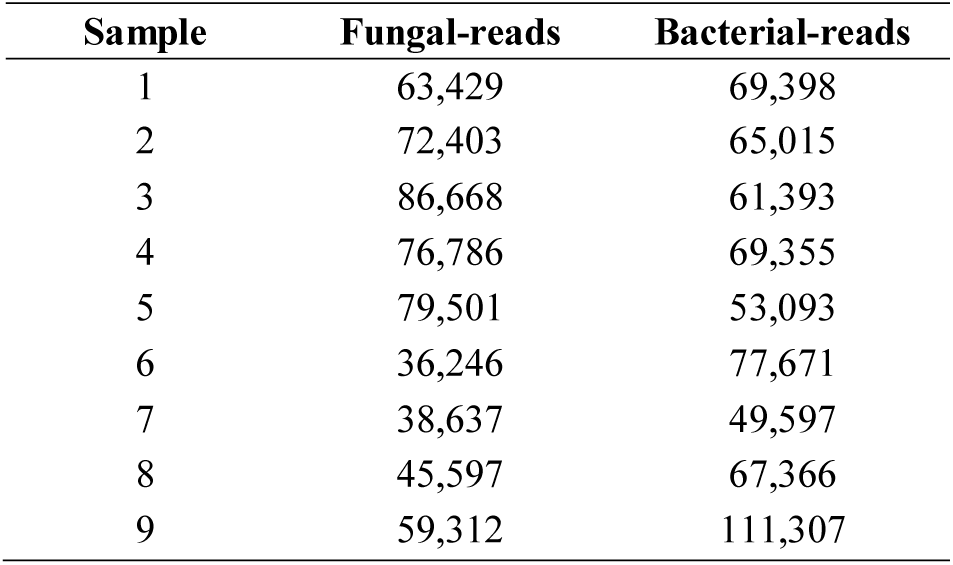
Number of Illumina Miseq reads allocated to each sample.

**Figure 1.**
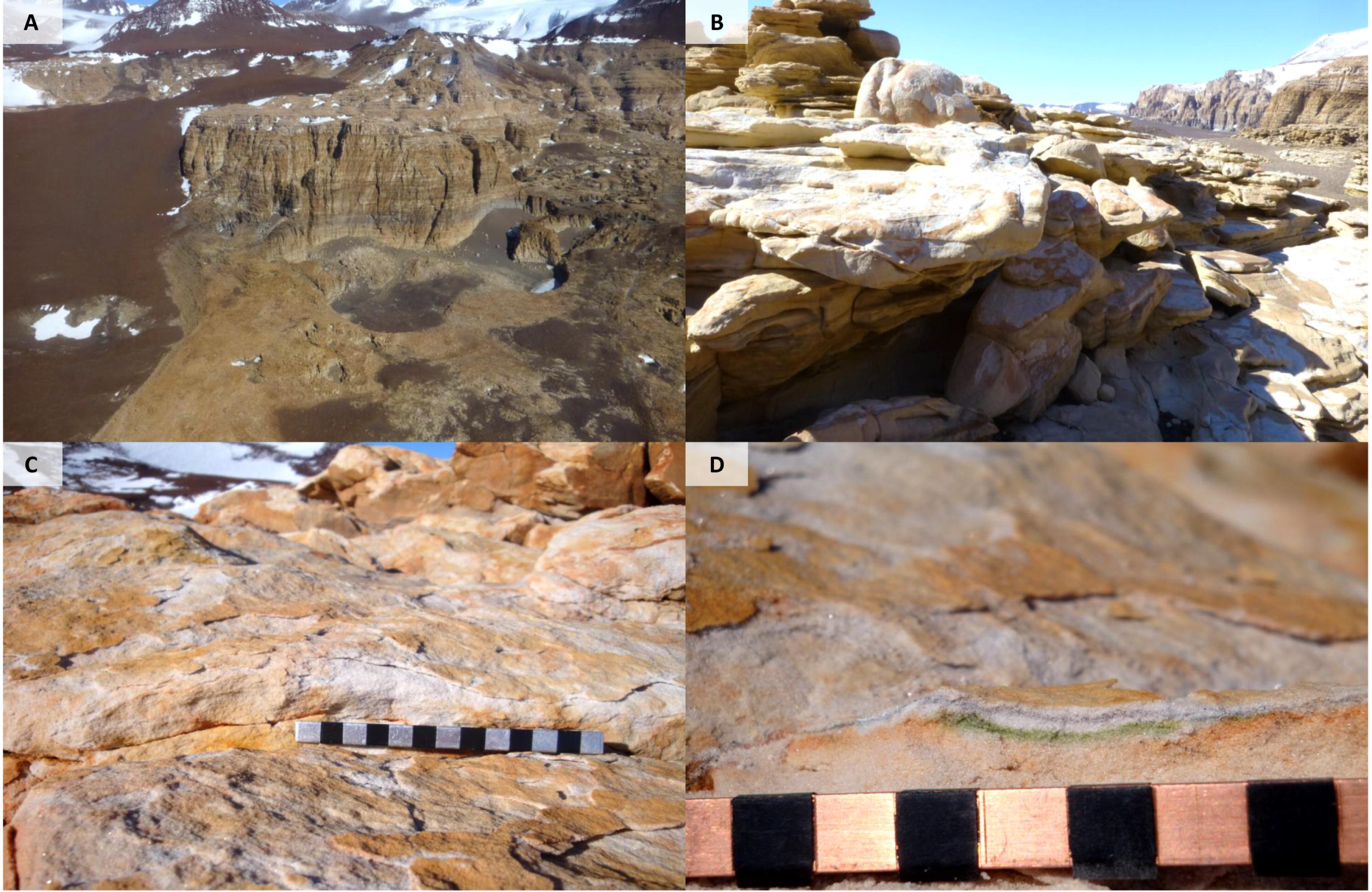
Landscape of Battleship Promontory (McMurdo Dry Valleys), visited during the XXVI Italian Antarctic Expedition (2010-2011) (A); Sandstone outcrops (B,C); Cryptoendolithic colonization just below the rock surface (D).

### Biodiversity and community composition

Species richness (S), Shannon (H’) and Simpson (1-D) indices were calculated for each rock sample. The observed fungal richness recorded ranged from 153 to 101 OTUs. In contrast, bacteria exhibited a higher diversity richness, ranging from 160 to 470 OTUs. Globally, the biodiversity, estimates from the 16S and ITS sequences, indicated that H’ ranged from 1.58 to 2.63 in fungi and 1.74 to 3. 70 bacteria; 1-D estimate of diversity ranged from 0.72 to 0.98 in ITS sequences and from 0.68 to 0.87 in 16S sequences (Table 3). Pairwise comparisons of biodiversity measures using Tukey’s test reported that, in generally, biodiversity varied significantly amongst samples and even, between fungi and bacteria (*p*<0.05, data not shown).

**Table 3.**
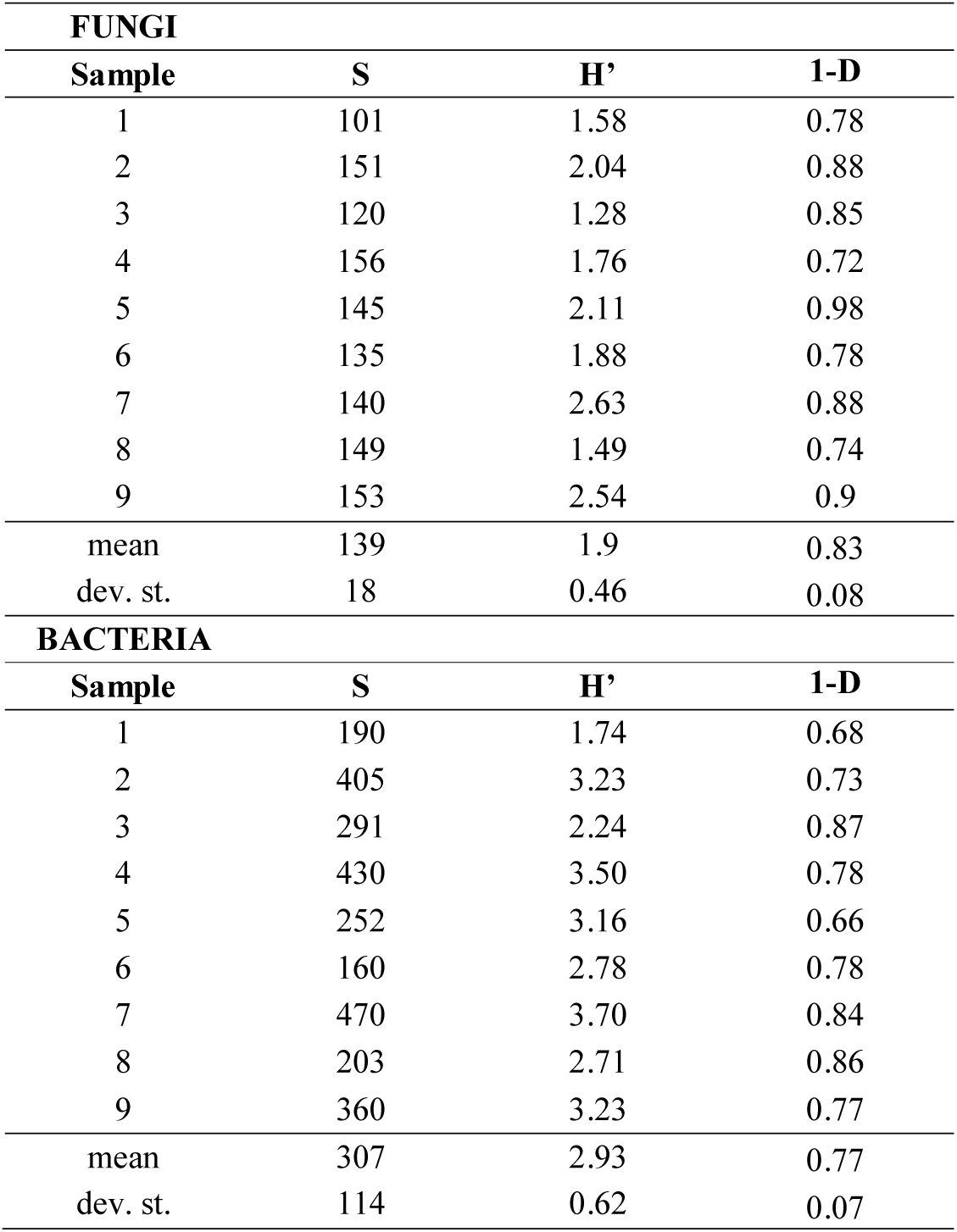
Diversity metrics for fungal and bacterial rRNA sequences for each sample. Species richness (S), Shannon index (H’) and Simpson’s Index of Dominance (1-D) are reported.

| FUNGI |  |  |  |
| --- | --- | --- | --- |
| Sample | S | H' | 1-D |
| 1 | 101 | 1.58 | 0.78 |
| 2 | 151 | 2.04 | 0.88 |
| 3 | 120 | 1.28 | 0.85 |
| 4 | 156 | 1.76 | 0.72 |
| 5 | 145 | 2.11 | 0.98 |
| 6 | 135 | 1.88 | 0.78 |
| 7 | 140 | 2.63 | 0.88 |
| 8 | 149 | 1.49 | 0.74 |
| 9 | 153 | 2.54 | 0.9 |
| mean | 139 | 1.9 | 0.83 |
| dev. st. | 18 | 0.46 | 0.08 |
| BACTERIA |  |  |  |
| Sample | S | H' | 1-D |
| 1 | 190 | 1.74 | 0.68 |
| 2 | 405 | 3.23 | 0.73 |
| 3 | 291 | 2.24 | 0.87 |
| 4 | 430 | 3.50 | 0.78 |
| 5 | 252 | 3.16 | 0.66 |
| 6 | 160 | 2.78 | 0.78 |
| 7 | 470 | 3.70 | 0.84 |
| 8 | 203 | 2.71 | 0.86 |
| 9 | 360 | 3.23 | 0.77 |
| mean | 307 | 2.93 | 0.77 |
| dev. st. | 114 | 0.62 | 0.07 |

A non-metric multidimensional scaling (NMDS) ordination was computed to investigate changes in communities composition using both incidence data (Jaccard dissimilarity index) and the read-abundance data (Bray–Curtis metrics), separately, to avoid the uncertainty whether read abundance was a good indicator of OTU abundance in the samples (Amend *et al*., 2010). Since both approaches produced similar outputs, results based on abundance data are reported only. NMDS plots resulted in two-dimensional solutions with stress values of 0.095 (fungi) and 0.076 (bacteria). Fungal assemblages had more similarity (p<0.05) among all samples than did the bacteria, where only few samples were similar (p>0.05) (Figure 2).

**Figure 2.**
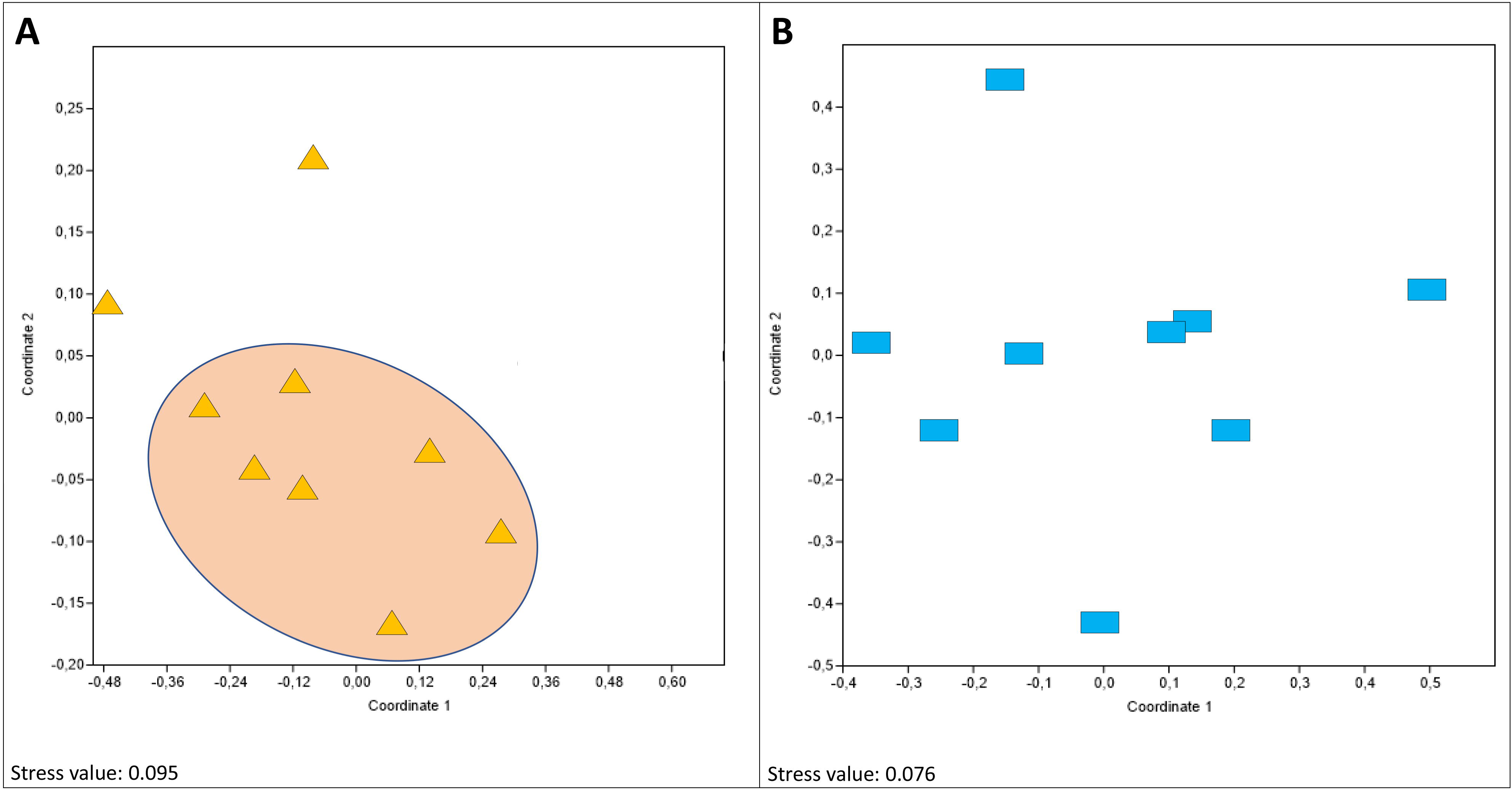
Non-Metric Multidemensional Scaling (N-MDS) ordination plots for fungal (A) and bacterial (B) groups of Antarctic endolithic communities collected at Battleship Promontory, based on square-root transformed abundance data.

### Estimating and determining sampling effort

The species accumulation curve is a graph of the expected number of detected species as a function of sampling e□ort, representing the sampling process (Palmer, 1990). Fungal randomized accumulation curve reached saturation with our sampling effort and the asymptote indicated that 100% of the species present were detected when nine samples are included (Figure 3A). In contrast, the total bacterial diversity was much higher and accumulation curves showed a continued increase in species per sample, indicating that biodiversity of the entire resident community was not captured and sampling was not exhaustive (Figure 3B). An extrapolation analysis of empirical sample-based rarefaction curves estimated the number of bacterial species expected to be found in a larger number of samples from the same assemblage. Figure 4 showed that, increasing the number of samples, the sampling would require approximately four times more effort (□40 samples) for a comprehensive description of bacterial diversity.

**Figure 3.**
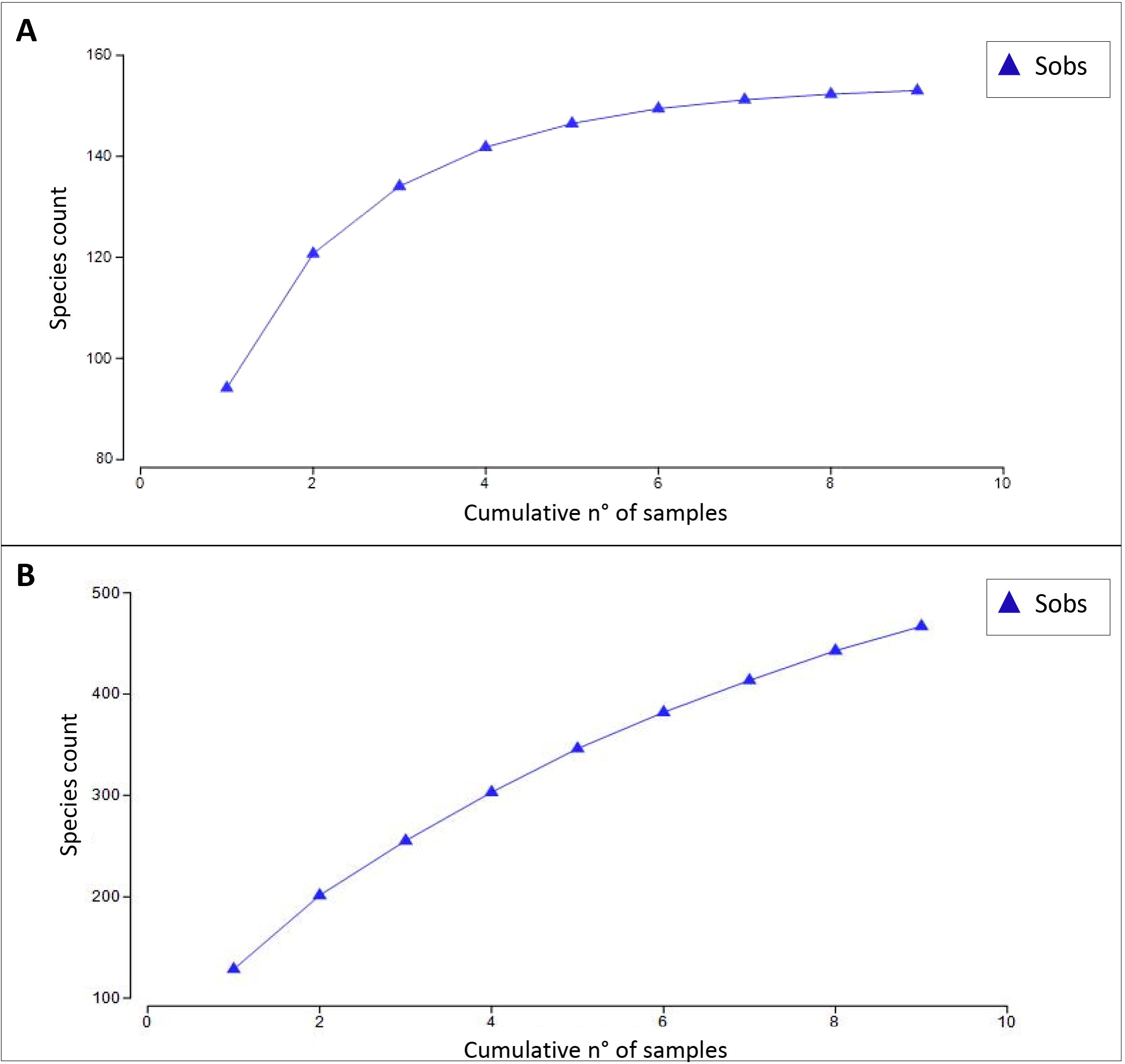
Fungal (A) and bacterial (B) species accumulation curve produced by the analytical formulae and randomizing (999 permutations) the samples via PRIMER-E software. Values were obtained using observed species (Sobs) counts and plotted against increasing number of samples.

**Figure 4.**
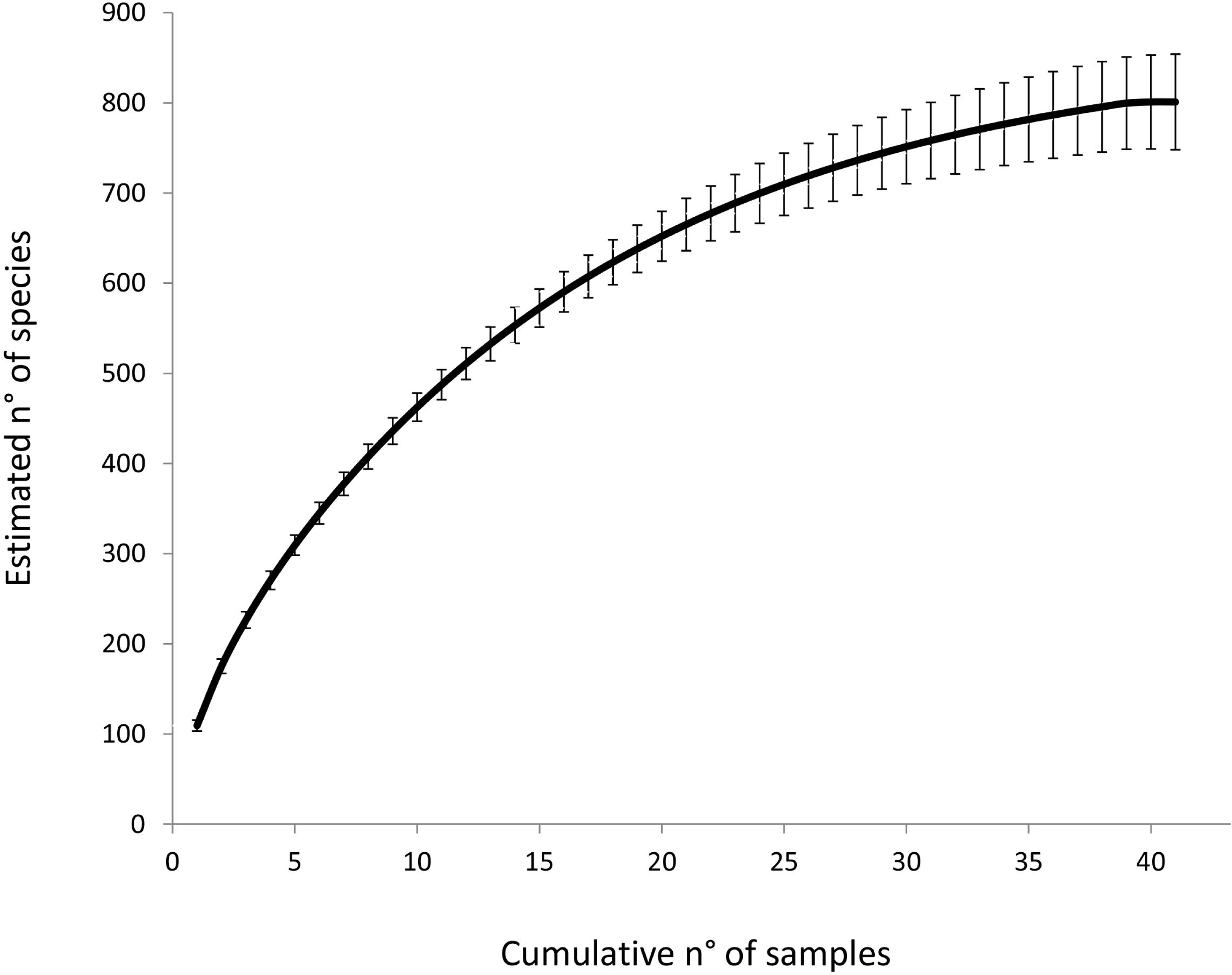
Expected species richness values (solid line) were computed for augmented reference samples (sample-based extrapolation curves) using EstimateS with 95% confidence intervals. Error bars represent the deviation standards.

### Fungal and bacterial abundance

The abundance of fungi and bacteria, as determined using qPCR, varied significantly (p<0.05) along whole sampling ranging from 5.3 × 10^5^ to 2.8 × 10^6^ fungal and from 5.8 × 10^4^ to 1.3 × 10^5^ bacterial gene copies, respectively (see Fig. 5 and Table 1S). Particularly, the lowest fungal abundance was estimated in sample 3 and 5 (527,384 and 822,113 copies, respectively), while the highest in sample 6 (2,784,953 copies); in bacterial compartment, 8 and 9 showed the lowest abundance (55,679 and 57,923 copies, respectively), while the highest value (128,796) was retrieved in sample 3 (Table 1S). Additionally, fungal and bacterial abundances significantly varied (p<0.05) along whole sampling; this resulted more remarkable in fungi, where almost all samples are different, while bacterial abundances were more homogenous.

Fungal-to bacterial ratio (F/B) (based on log copy numbers) was also calculated from these qPCR results. These findings showed a slightly higher dominance of fungi in these ecosystems varying between 1.25 ± 0.06 (mean ± SD).

**Figure 5.**
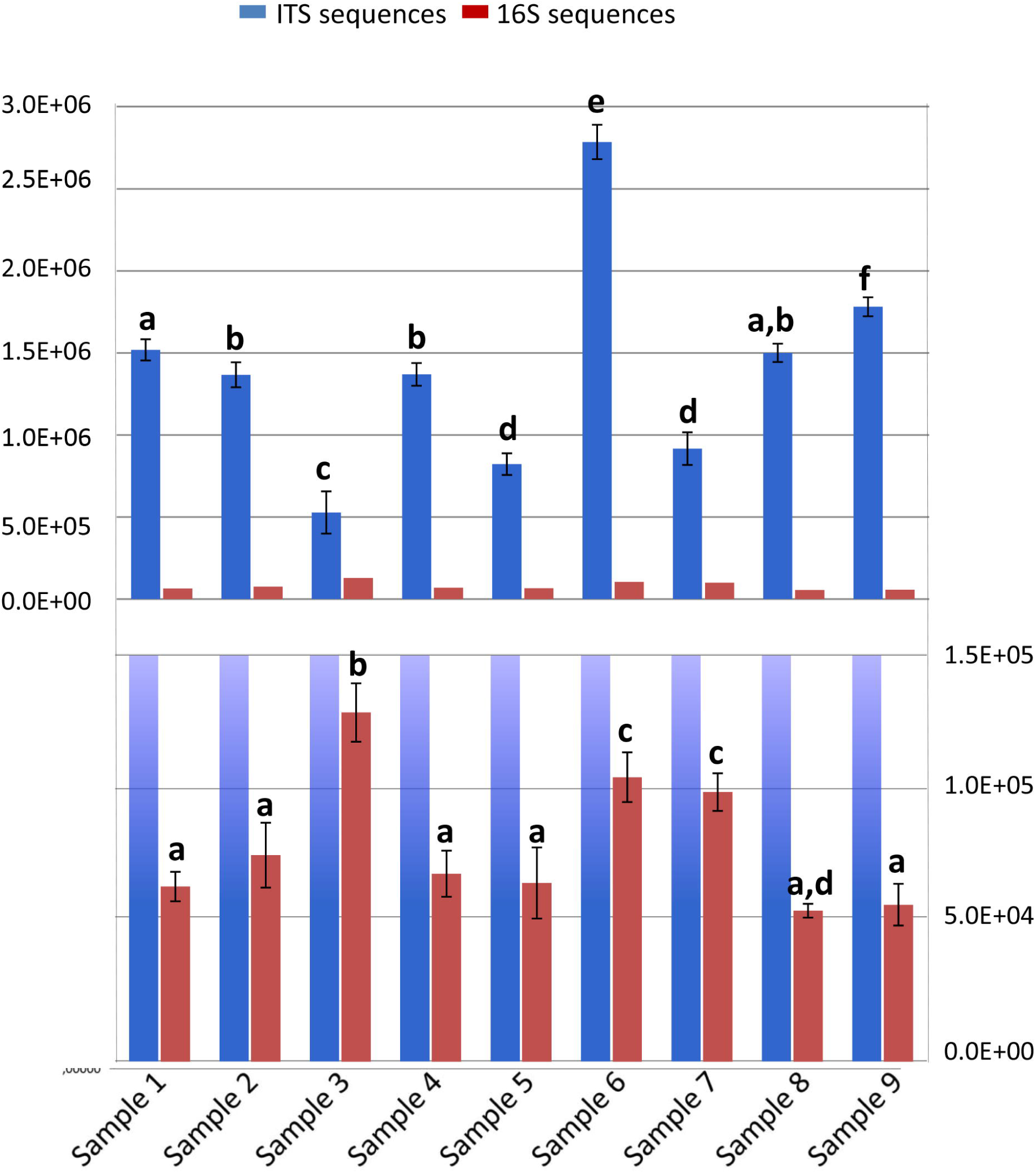
The abundance of fungi (**a**) and bacteria (**b**), as indicated by the number of ITS or 16S ribosomal DNA (rDNA) copies measured using quantitative PCR (qPCR) assay. Error bars are the standard errors. Significant differences, calculated by Tukey test (p<0.05), are indicated by different letters.

## Discussion

The McMurdo Dry Valleys in Southern Victoria Land, is one of most extreme ice-free areas known on Earth. The extreme aridity, temperature swings, and UV radiation exposure have been used to classify it as the closest terrestrial analogue of Martian environment on Earth. In this region, where climatic conditions are too harsh to allow colonization on the surface of rocks, endolithic communities are almost the only life-form possible. Rock niches provides buffered conditions for microorganisms to find a refuge for survival (Friedmann and Ocampo-Friedmann, 1976; Nienow and Friedmann, 1993).

Recent molecular advances have provided new insights into microbial diversity and community composition in Antarctic cryptoendolithic communities. These ecosystems harbour a very low diversity of both eukaryotic and prokaryotic assemblages, dominated by a restricted number of highly specialized species (Archer et al., 2017; Selbmann et al. 2017; Coleine et al., 2018a, b). To better define the minimum sampling effort needed to capture the community diversity and to inform and optimize future sampling protocols we evaluated how well our sampling recovered whole metacommunity composition from Antarctic cryptoendolithic communities.

The high degree of specialization shaped by the presence of very few predominant species and rare taxa as determined by the Shannon’s and Simpson’s dominance indices (0.83±0.08 and 0.77±0.0.7 in fungi and bacteria, respectively), suggested that these microbial communities are highly specialized, adapted and resistant, but they may be dramatically influenced by any disturbance events which may irreparably affect their fragile equilibrium and future in the face of global change (Selbmann et al., 2017; Coleine et al., 2018a, b). Analysis of these communities and correlations to abiotic and biotic parameters will help determine and predict the responses of ecosystem processes to climate change.

Overall, a considerable amount of heterogeneity has been observed between rock samples, both for fungal and bacterial assemblages; the high degree of variability may be related to the breadth of the sampling area (100 m^2^). A more homogeneous dataset may be observed from a selection of rocks collected in strict proximity to each other. A remarkable variability in biodiversity and structure in endolithic communities collected from the same location was reported by Selbmann and colleagues (2017) or in sandstones collected in localities with similar environmental parameters (altitude and sea distance) as recently reported by Coleine et al. (2018a). Specifically, in our survey, richness and Shannon’s biodiversity index values significantly varied among all samples: number of observed species ranged from 93 to 153 and 160 to 470 in fungi and bacteria, respectively, while biodiversity values from 1.58 to 2.63 and 1.74 to 3.70 in fungi and bacteria, respectively. The higher values of bacteria are consistent with a reconstruction based on a tree of life that found bacteria generally more diverse than fungi (Hinchliff et al., 2015); richness in species in individual communities varied according to the habitat, but it seems to be definitively higher for bacteria than fungi. For example, in soil communities, bacterial species are generally 2–3 times more numerous than fungi (Urbanová et al., 2015).

Changes in community’s composition was reported both in fungi and bacteria, as suggested by NMDS ordinations, even in samples with comparable species richness. Moreover, community composition was considerably more different among observations of bacteria in the samples. A PERMANOVA analysis showed that community structure varied in almost all samples (p>0.05) and these data were consistent when analyzed as presence-absence or abundance matrices.

Substantial heterogeneity was even observed considering fungal and bacterial relative abundances estimated with qPCR assay. Abundance data was significantly variable along the sampled area (values ranged 5.3 × 105 to 2.8 × 106 for fungi and from 5.8 × 104 to 1.3 × 105 for bacteria). The fungal/bacterial (F/B) ratio, a common metric to assess environmental impacts and the functional implications of microbial communities (Raeymaekers, 2000; Fierer et al., 2005), was also assayed in order to compare community makeup. We found a weak fungal dominance (F/B: 1.25); this may be explained by the ability of fungi to grow at lower water activity than prokaryotes, the accuracy of F/B ratio from Antarctic endolithic communities remains unexplored; future research into the microbial members of these environments is needed, especially gathering a basic understanding of the ecology of bacteria and fungi in these ecosystems in order to better assess the mechanism by which a change in F/B dominance might occur. For example, in acidic soils the fungi are predominant biomass linked to the observation that fungi are weakly influenced by pH values (Rousk et al., 2010), while bacterial biomass is highly affected by changes in pH (Griffiths et al., 2011). Soil nutrient availability and moisture may also affect the F/B ratio (Strickland and Rousk, 2010).

Based on these observations, properly describing the complete microbial biodiversity in Antarctic cryptoendolithic communities remains a challenge. The high variability in microbial assemblages in the same sampled area suggests that there are highly localized patterns in individual rocks. Species accumulation curves indicated that fungal diversity in the sampled area may be sufficiently captured with the current sampling depth and design (nine sandstone samples). The number of species found exhibited a cumulative pattern of diminishing increase with increasing the samples number. Conversely, we captured only a subset of the total estimated bacterial diversity based on the lack asymptote of the accumulation curves. We estimate a total of 40 samples from nine rocks is needed to reach a plateau in the observations of bacteria community diversity.

This work emphasizes the importance of sampling efforts on precision and provides information to determine sampling efficiency in extreme areas like the Antarctic where extensive sampling can be prohibitive. Based on these results, we determined that the prior sampling effort had was able to robustly assess species richness and community composition in Antarctic cryptoendolithic ecosystems, especially for the fungal component. Recommendations for future eDNA metabarcoding surveys and research would need to take into account these expected scale of local biodiversity within rocks in order to fully assess cryptoendolithic community diversity. Since a large number of additional undetected bacterial species are likely to be present in the sampled area, for future studies we suggest a pooling strategy, prior to DNA extraction, to minimize local variability. The pooling method in environmental metabarcoding studies may be particularly useful for among-site comparison of soil and rock communities and it is generally utilized to reduce sample numbers for evaluations of microbial community structure (Baker et al., 2009; Osborne et al., 2011). Typically pooling of eDNA is not recommended since DNA of rare species could be diluted and important portion of microbial composition may be under sampled, especially if communities are extremely complex (Manter et al., 2010). These approaches can be considered due to the very low biodiversity and complexity of endolithic ecosystems inhabiting ice-free area of Victoria Land in Antarctica (Selbmann et al., 2017; Coleine et al., 2018a).

This pilot study sought to define optimal sampling efforts for future studies and further investigations will refine hypotheses regarding local distribution patterns of microbes. The current sampling of areas of Southern Victoria Land can be contrasted to environmentally different Northern Victoria Land. This examination of diversity using eDNA methods gives insights on optimizing sampling protocols to increase the sampling efficiency and decrease an unnecessary number of samples which can reduce the impact on delicate ecosystems and strictly protected areas.

### Compliance with Ethical Standards

This article does not contain any studies with human participants or animals performed by any of the authors. Conflict of Interest: The authors declare that they have no conflict of interest. The datasets generated during the current study are available in the GenBank repository under BioProject accession number PRJNA379160.

## ACKNOWLEDGMENTS

L.S., C.C. and L.Z. wish to thank the Italian National Program for Antarctic Researches (PNRA) for funding sampling campaigns and researches activities in Italy in the frame of 2009/A1.11, 2013/AZ-17, 2015/AZ1.02 and AMunDsEN PNRA_00006 projects. The Italian Antarctic National Museum (MNA) is acknowledged for financial support to the Mycological Section of the MNA for preserving rock Antarctic samples analysed in this study and stored in the Culture Collection of Fungi from Extreme Environments (CCFEE), University of Tuscia, Italy.

Fungal ITS and bacterial 16S primers were made available through the Alfred P. Sloan Foundation Built Environment Program and sequencing supported by funds through United States Department of Agriculture - National Institute of Food and Agriculture Hatch project CA-R-PPA-5062-H to JES. N.P. was supported by a Royal Thai government fellowship. Data analyses were performed on the High-Performance Computing Cluster at the University of California-Riverside in the Institute of Integrative Genome Biology supported by NSF DBI-1429826 and NIH S10-OD016290.

---

**Figure 1S.** The frequency of observed OTUs for each sample was used to calculate fungal rarefaction curves using the ‘rarecurve’ function in the R library vegan v2.3-4 (Oksanen *et al*., 2013) in R 3.2.0 (R-Development-Core-Team 2015).

**Figure 2S.** The frequency of observed OTUs for each sample was used to calculate bacterial rarefaction curves using the ‘rarecurve’ function in the R library vegan v2.3-4 (Oksanen *et al*., 2013) in R 3.2.0 (R-Development-Core-Team 2015).

**Table 1S.** The abundance of fungi and bacteria, as indicated by the number of ITS and 16S ribosomal DNA (rDNA) copies measured using quantitative PCR (qPCR) assay.

**Table 2S.** Fungal taxonomy results at 97% of identity. Taxonomy assignment was performed with SINTAX/UTAX databases (Edgar, 2010).

**Table 3S.** Bacterial taxonomy results at 97% of identity. Taxonomy assignment was performed with SINTAX/UTAX databases (Edgar, 2010).

