## Supplementary figures and images for "Sampling strategies to assess microbial diversity of Antarctic cryptoendolithic communities"

### Figure 1S

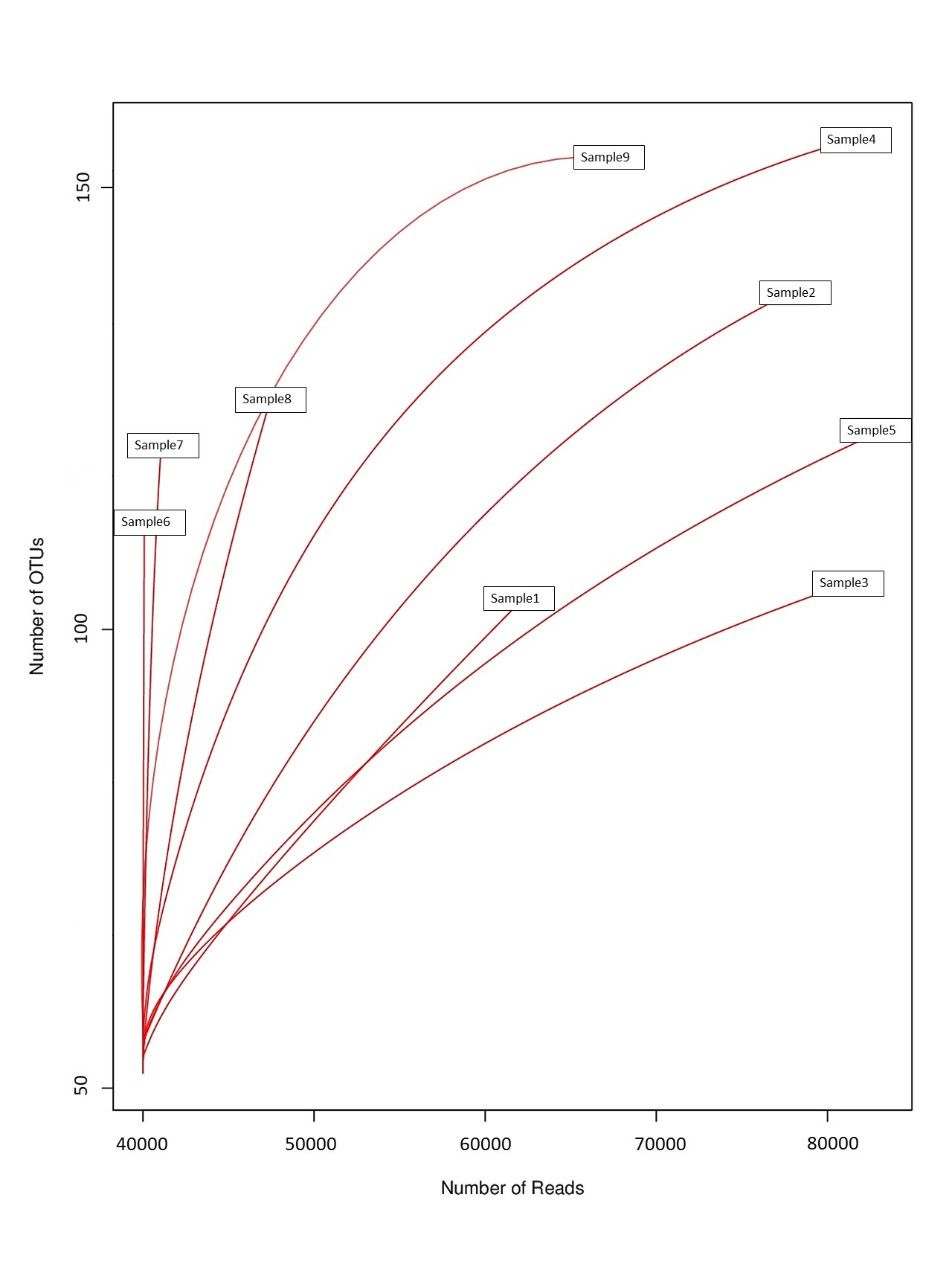

### Figure 2S

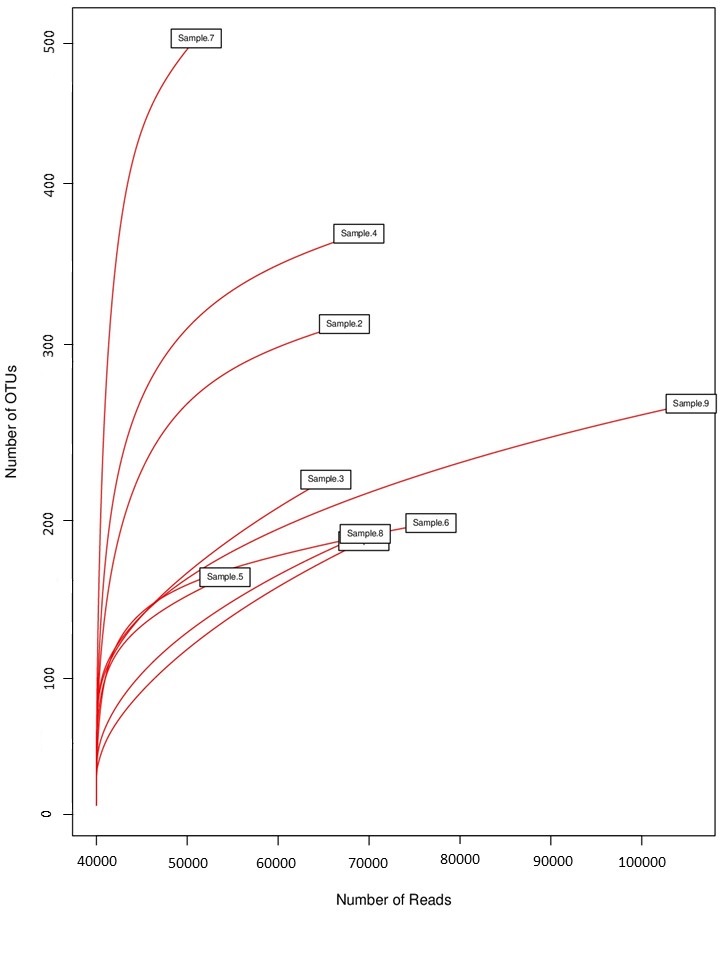
